# Object-in-Place Memory Predicted by Anterolateral Entorhinal Cortex and Parahippocampal Cortex Volume in Older Adults

**DOI:** 10.1101/409607

**Authors:** Lok-Kin Yeung, Rosanna K. Olsen, Bryan Hong, Valentina Mihajlovic, Maria C. D’Angelo, Arber Kacollja, Jennifer D. Ryan, Morgan D. Barense

## Abstract

The lateral portion of the entorhinal cortex is one of the first brain regions affected by tau pathology, an important biomarker for Alzheimer’s disease (AD). Improving our understanding of this region’s cognitive role may help identify better cognitive tests for early detection of AD. Based on its functional connections, we tested the idea that the human anterolateral entorhinal cortex (alERC) may play a role in integrating spatial information into object representations. We recently demonstrated that the volume of the alERC was related to processing the spatial relationships of the features *within* an object (Yeung et al., 2017). In the present study, we investigated whether the human alERC might also play a role in processing the spatial relationships *between* an object and its environment using an eyetracking task that assessed visual fixations to a critical object within a scene. Guided by rodent work, we measured both object-in-place memory, the association of an object with a given context (Wilson et al., 2013), and object-trace memory, the memory for the former location of objects (Tsao, Moser, & Moser, 2013). In a group of older adults with varying stages of brain atrophy and cognitive decline, we found that the volume of the alERC and the volume of the parahippocampal cortex (PHC) selectively predicted object-in-place memory, but not object-trace memory. These results provide support for the notion that the alERC may integrate spatial information into object representations.

## Introduction

Lateral portions of the entorhinal cortex are among the earliest regions to develop tau pathology, a key biomarker for Alzheimer’s disease (AD) (Braak & Braak, 1991; Khan et al., 2014). In turn, the presence of tau pathology here is strongly related to local grey matter loss (Maass et al., 2017; Sepulcre et al., 2016). Consistent with these findings, recent work from our group showed smaller anterolateral entorhinal cortex (alERC) volumes in ostensibly-healthy older adults demonstrating early signs of preclinical AD-related cognitive decline (Olsen, Yeung et al., 2017). An important challenge in AD research is that neurodegenerative changes occur years before cognitive deficits become apparent with standard neuropsychological assessments (Sperling et al., 2011). Thus, finding a subtle cognitive effect specifically related to alERC neurodegeneration could significantly improve early detection of AD. However, these efforts are limited by a lack of understanding regarding the cognitive role of human entorhinal cortex subdivisions.

In rodents and nonhuman primates, it is well established that distinct subregions of the entorhinal cortex mediate two input pathways into the hippocampus. One pathway, thought to emphasize the processing of object information, originates in the ventral visual stream and projects to the lateral entorhinal cortex (LEC) via the perirhinal cortex (PRC) (Naber, Caballero-Bleda, Jorritsma-Byham, & Witter, 1997; Suzuki & Amaral, 1994; see also Cowell, Bussey, & Saksida, 2010). The other pathway, thought to emphasize the processing of spatial or contextual information, originates in the dorsal visual stream and projects primarily to the medial entorhinal cortex (MEC) via the postrhinal/parahippocampal cortex (PHC) (Burwell, 2000; Moser, Kropff, & Moser, 2008). The functional organization that arises from this anatomical organization has been described in two models. The PMAT (posterior medial, anterior temporal) model proposes that these two pathways form part of two distinct systems, supporting item and spatial/contextual information respectively (Ritchey, Libby, & Ranganath, 2015). The representational-hierarchical model proposes a hierarchical organization of stimulus representations of increasing complexity moving forward in each pathway (Cowell et al., 2010). Both models highlight the role of the hippocampus as where these two kinds of information, from the two entorhinal cortex subfields, converge.

Despite the extensive research on these two pathways in nonhuman animals, far less is known about their function and organization in humans. Recent work suggests there exists a similar functional parcellation of the human entorhinal cortex (Maass, Berron, Libby, Ranganath, & Düzel, 2015; Navarro Schröder, Haak, Zaragoza Jimenez, Beckmann, & Doeller, 2015). More specifically, functional connectivity analyses reveal that the human entorhinal cortex can be divided into two parts: an anterior-lateral subregion (the alERC) that co-activates with the PRC, and a posterior-medial subregion (the pmERC) that co-activates with the PHC. This suggests a functional homology between the human alERC with the rodent LEC, and between the human pmERC with the rodent MEC. Mirroring the functional distinctions proposed in the animal work, the PMAT model proposes that the PRC-alERC pathway is critical for representing object information and the PHC-pmERC pathway critical for representing spatial and contextual information (Ritchey et al., 2015). Functional neuroimaging data in humans supports this model, with greater BOLD activity in lateral ERC when processing the identity of a face or object, and greater BOLD activity in medial ERC when processing the spatial location of that object (Berron et al., 2018; Reagh et al., 2018; Reagh & Yassa, 2014; Schultz, Sommer, & Peters, 2012)

In the face of this apparent functional subdivision of the ERC, rodent studies have also reported that the separation between these two pathways is not absolute. Although the majority of connections continue along their respective pathways, the rodent LEC also has some reciprocal connections with the MEC (van Strien, Cappaert, & Witter, 2009). By analogy, similar connections between the homologous human alERC and pmERC might also exist. Based on these reciprocal connections between the two pathways, we and others have speculated that the alERC might play a role in integrating spatial information from the pmERC into the object representations supported by the PRC (Connor & Knierim, 2017; Yeung et al., 2017). Two LEC rodent studies provide support for this notion. Lesions to the rodent LEC led to impairments in “object-in-place” memory (i.e., memory for the association between an object and a spatial context), but not for memory for objects or spatial contexts independently (Wilson et al., 2013). Moreover, direct recording of the rodent LEC reported the presence of “object-trace cells”: place cells that fired specifically at the locations that had previously contained a certain object (Tsao et al., 2013). In humans, we recently found that alERC volume was positively related to processing the spatial relationship of features within an object (i.e., visual fixations to the configurally-relevant region of an object) (Yeung et al., 2017). In the present study, we sought to investigate whether the human alERC might also play a broader role in processing spatial information about an object, beyond the within-object processing that we and others have observed (Berron et al., 2018; Reagh et al., 2018; Reagh & Yassa, 2014; Schultz et al., 2012; Yeung et al., 2017). In particular, we assessed whether the integrity of the alERC was related to associating an object with its spatial context, as has been observed in rodents.

Inspired by the rodent work, we leveraged an eyetracking-based behavioral paradigm to test whether the human alERC, and surrounding medial temporal lobe (MTL) regions, might also play a role in object-in-place memory (i.e. associating an object with a particular location in a particular context) and/or object-trace memory (i.e. memory for the previous location of an object) (Ryan, Althoff, Whitlow, & Cohen, 2000; Ryan, Leung, Turk-Browne, & Hasher, 2007; Ryan & Cohen, 2004b, 2004a; Smith, Hopkins, & Squire, 2006; Smith & Squire, 2008). As in our previous work, we used eyetracking-based metrics as our outcome measures, as these measures are sensitive to memory effects which may not reach the level of conscious awareness (Ryan et al., 2000) and allowed us to more closely match our design to the aforementioned rodent studies. A group of older adult participants with varying levels of cognitive decline incidentally viewed computer-generated scenes that were either entirely novel, repeated identically from the previous viewing, or were manipulated such that a single critical object was moved. This allowed us to derive eyetracking-based measures of object-in-place and object-trace memory, based on fixations to the location currently or previously occupied by the object in the manipulated scenes, respectively. The novel scenes were used to assess global measures of novelty detection. Further, we employed a recently-developed manual segmentation protocol to assess the volume of the alERC (Maass et al., 2015; Olsen, Yeung et al., 2017) and surrounding hippocampal subfields and MTL cortices (Olsen et al., 2013). We hypothesized that the volumes of the alERC, along with the PHC, pmERC and hippocampus, which belong to the spatial/contextual processing pathway, would relate to eyetracking-based measures of both object-in-place memory and object-trace memory.

## Methods

### Participants

Thirty-two community-dwelling older adults were recruited from the community in Toronto. Data from two participants were excluded due to eyetracker failure. The remaining thirty participants had a mean age of 72.3 years (SD: 5.2, range: 59-81, 23 women). Participants had previously been tested on the Montreal Cognitive Assessment (MoCA) (Nasreddine et al., 2005) within the last twenty-three months (mean: 12.1, SD: 6.8, range: 2-23), and were selected to provide a distribution of MoCA scores (mean: 25.3, SD: 3.0, range: 17-30). Given that MoCA is sensitive to the presence of mild cognitive impairment, which is associated with MTL/hippocampus volume loss (Jack et al., 1997), our intention was to select for a participant group that had a good distribution of cognitive abilities and MTL/hippocampal regional volumes. These participants were the subset of an original sample of forty participants from Olsen, Yeung et al., (2017) whom we were able to recruit for the present study (i.e. 8 participants were lost to follow-up). The original group of forty participants were chosen such that twenty participants had scored above the recommended MoCA cutoff score (≥26) and twenty participants had scored below the MoCA cutoff score (<25) (data on these participants has previously been reported in Olsen, Yeung et al., 2017 and Yeung et al., 2017). Of the thirty participants whose data we report here, fourteen scored above the MoCA cutoff score, and sixteen scored below it. These two groups were matched for age (participants in this study: t(28) = 1.29, p = 0.21, d = 0.237) and years of education (participants in this study: t(28) = 0.51, p = 0.61, d = 0.076). Due to our efforts to match participants above and below the MoCA cutoff score in terms of demographic characteristics, MoCA and age were not correlated among the thirty participants in this study (r = -0.250, p = 0.13). For the purposes of the present study, we were primarily interested in how MTL volume differences related to cognitive performance, rather than how participants who scored above/below the MoCA threshold differed; thus, we treated all the participants as a single group for all subsequent analyses. Participants received a battery of neuropsychological tests to characterize their cognitive status (Table 1) in an earlier session (mean interval = 10.2 months, SD = 8.8 months). All participants had normal or corrected-to-normal vision (with glasses or bifocals), and were screened for color blindness, psychological or neurological disorders, brain damage (i.e. stroke or surgery) and metal implants which would have precluded MR imaging. All participants gave informed consent. This research received ethical approvals from the Research Ethics Boards of the University of Toronto and Baycrest.

**Table 1:** Neuropsychological battery results expressed as means (SD). Maximum and cut-off scores for tests indicated in parentheses in left column. WMS-IV = Wechsler Memory Scale, 4^th^ Ed. Note that two participants did not complete the subjective memory questionnaire

| Test | All participants (N=32) | Participants included in data analysis (N=30) |
| --- | --- | --- |
| MoCA (/30) | 25.3 (2.9)<br>Slightly impaired | 25.3 (2.9)<br>Slightly impaired |
| WMS-IV Logical Memory |  |  |
| Immediate Recall Scaled Score (/20) | 11.3 (2.8)<br>63.7%ile | 11.4 (2.9)<br>64.2%ile |
| Delayed Recall Scaled Score (/20) | 10.8 (2.7)<br>59.1%ile | 10.9 (2.8)<br>59.3%ile |
| Recognition Accuracy | 78.7% (17.8%) | 81.8% (11.1%) |
| Trails A | 43.9s (14.1s)<br>40.9%ile | 44.1s (14.4s)<br>40.6%ile |
| Trails B | 98.90 (35.3s)<br>52.0%ile | 95.5s (34.5s)<br>53.2%ile |
| Digit Span Forward Score (/16) | 10.1 (2.3)<br>50.6%ile | 10.2 (2.3)<br>51.9%ile |
| Digit Span Backward Score (/14) | 6.3 (2.3)<br>26.9%ile | 6.4 (2.3)<br>28.5%ile |
| Rey-Osterrieth Complex Figure |  |  |
| Copy (/32) | 26.8 (5.7)<br>27.9%ile | 26.6 (5.8)<br>27.6%ile |
| Immediate Recall (/32) | 11.7 (6.5)<br>37.2%ile | 12.1 (6.4)<br>39.5%ile |
| Delayed Recall (/32) | 10.4 (6.6)<br>32.6%ile | 10.7 (6.7)<br>34.3%ile |
| Weschler Abbreviated Scale of Intelligence |  |  |
| Vocabulary (/80) | 58.4 (10.1)<br>63.9%ile | 58.7 (10.1)<br>64.8%ile |
| Similarities (/48) | 35.8 (5.2)<br>74.0%ile | 36.0 (5.2)<br>74.9%ile |
| Matrix Reasoning (/32) | 20.7 (6.8)<br>72.2%ile | 20.6 (7.0)<br>71.5%ile |
| Block Design (/71) | 28.8 (14.7)<br>52.9%ile | 29.3 (14.9)<br>53.6%ile |
| Visual Object and Spatial Perception Battery |  |  |
| Shape Detection (/20)<br>(Cut-off score < 15) | 19.1 (1.2)<br>Pass | 19.0 (1.2)<br>Pass |
| Incomplete Letters (/20)<br>(Cut-off score < 16) | 19.2 (0.9)<br>Pass | 19.2 (0.9)<br>Pass |
| Dot Counting (/10)<br>(Cut-off score < 8) | 9.8 (0.4)<br>Pass | 9.8 (0.4)<br>Pass |
| Position Discrimination (/20)<br>(Cut-off score < 18) | 19.1 (1.7)<br>Pass | 19.1 (1.7)<br>Pass |
| Number Location (/10)<br>(Cut-off score < 7) | 9.0 (1.7)<br>Pass | 9.0 (1.7)<br>Pass |
| Cube Analysis (/10)<br>(Cut-off score < 6) | 9.4 (1.3)<br>Pass | 9.3 (1.3)<br>Pass |
| Silhouettes (/30)<br>(Cut-off score < 15) | 19.3 (5.3)<br>Pass | 19.2 (5.4)<br>Pass |
| Object Decision (/20)<br>(Cut-off score < 14) | 16.6 (2.0)<br>Pass | 16.7 (2.0)<br>Pass |
| Progressive Silhouettes (/20)<br>(Cut-off score > 15) | 10.4 (3.3)<br>Pass | 10.2 (3.2)<br>Pass |
| Subjective memory rating<br>(Memory functioning<br>questionnaire, /448) | 290.6 (54.2)<br>Minimal subjective<br>difficulties | 291.9 (55.3)<br>Minimal subjective<br>difficulties |

### MRI Scan Parameters

High-resolution T2-weighted images were acquired in an oblique-coronal plane, perpendicular to the long axis of the hippocampus (TE/TR = 68ms/3000ms, 20-28 slices depending on head size, 512x512 acquisition matrix, voxel size = 0.43 x 0.43 x 3 mm, no skip, FOV = 220mm), on a 3T Siemens Trio scanner at the Rotman Research Institute at Baycrest (Toronto, ON). The first slice was placed anterior to the appearance of the collateral sulcus (including the temporal pole where possible), and the last slice placed posterior to the hippocampal tail to ensure full coverage of the entire hippocampus and all of the MTL cortices included in the volumetric analyses for all participants. To confirm slice placement, a T1-weighted MP-RAGE whole brain anatomical scan (TE/TR = 2.63 ms/2000ms, 176 slices perpendicular to the AC-PC line, 256x192 acquisition matrix, voxel size = 1 x 1 x 1 mm, FOV = 256mm) was acquired immediately prior to the T2-weighted scan. The T1-weighted images were also used to estimate total intracranial volume for head-size correction (see volume correction section below).

### Manual Segmentation

For each participant, L.Y. manually segmented three hippocampal subfields (CA1, dentate gyrus/CA2&3, and subiculum) and four MTL cortices (anterolateral entorhinal cortex (alERC), posteromedial entorhinal cortex (pmERC), perirhinal cortex (PRC), and parahippocampal cortex (PHC)) on coronal slices of the T2-wighted structural scans (in-plane resolution: 0.43x0.43mm, 3mm between slices) using FSLview (v3.1). Manual segmentation followed the Olsen-Amaral-Palombo (OAP) protocol (Olsen et al., 2013; Palombo et al., 2013; see also the appendix to Yushkevich et al., 2015a) supplemented with a modified version of the protocol provided by Maass et al. (2015) for the subdivisions of the entorhinal cortex – see Figure 1 for a visualization of the segmentation protocol. Average volumes for each manually segmented brain region are presented in Table 3, and correlations between brain region volumes are presented in Table 4.

**Table 3:** Average volumes (±SD, in mm^3^) for each of the three manually-segmented hippocampal subfields and four MTL cortices segmented for this study (corrected for head-size)

| Brain Region |  | All participants (N=32) | Participants included in data analysis (N=30) |
| --- | --- | --- | --- |
| Hippocampus | CA1 | 1238.41 ± 149.33 | 1237.15 ± 154.22 |
| | Subiculum | 1105.20 ± 192.91 | 1103.46 ± 194.80 |
| | DG/CA23 | 1957.19 ± 351.09 | 1963.32 ± 357.11 |
| MTL | PRC | 4966.35 ± 1154.23 | 4954.03 ± 1118.44 |
|  | alERC | 1343.32 ± 274.72 | 1330.96 ± 278.97 |
|  | pmERC | 440.84 ± 112.80 | 443.50 ± 113.33 |
|  | PHC | 3640.21 ± 636.00 | 3649.43 ± 655.53 |

**Table 4:** Pearson’s correlations between volumes of manually segmented brain regions; * p < 0.05, ** p < 0.01.

|  | CA1 | Subiculum | DG/CA23 | PRC | alERC | pmERC | PHC |
| --- | --- | --- | --- | --- | --- | --- | --- |
| CA1 | 1 | .375* | .671** | .366* | .318* | .347* | .440** |
| Subiculum |  | 1 | .162 | -.272 | -.015 | .590** | .228 |
| DG/CA23 |  |  | 1 | .468** | .258 | .219 | .311 |
| PRC |  |  |  | 1 | .395* | .011 | .159 |
| alERC |  |  |  |  | 1 | .198 | .201 |
| pmERC |  |  |  |  |  | 1 | .274 |
| PHC |  |  |  |  |  |  | 1 |

**Figure 1:**
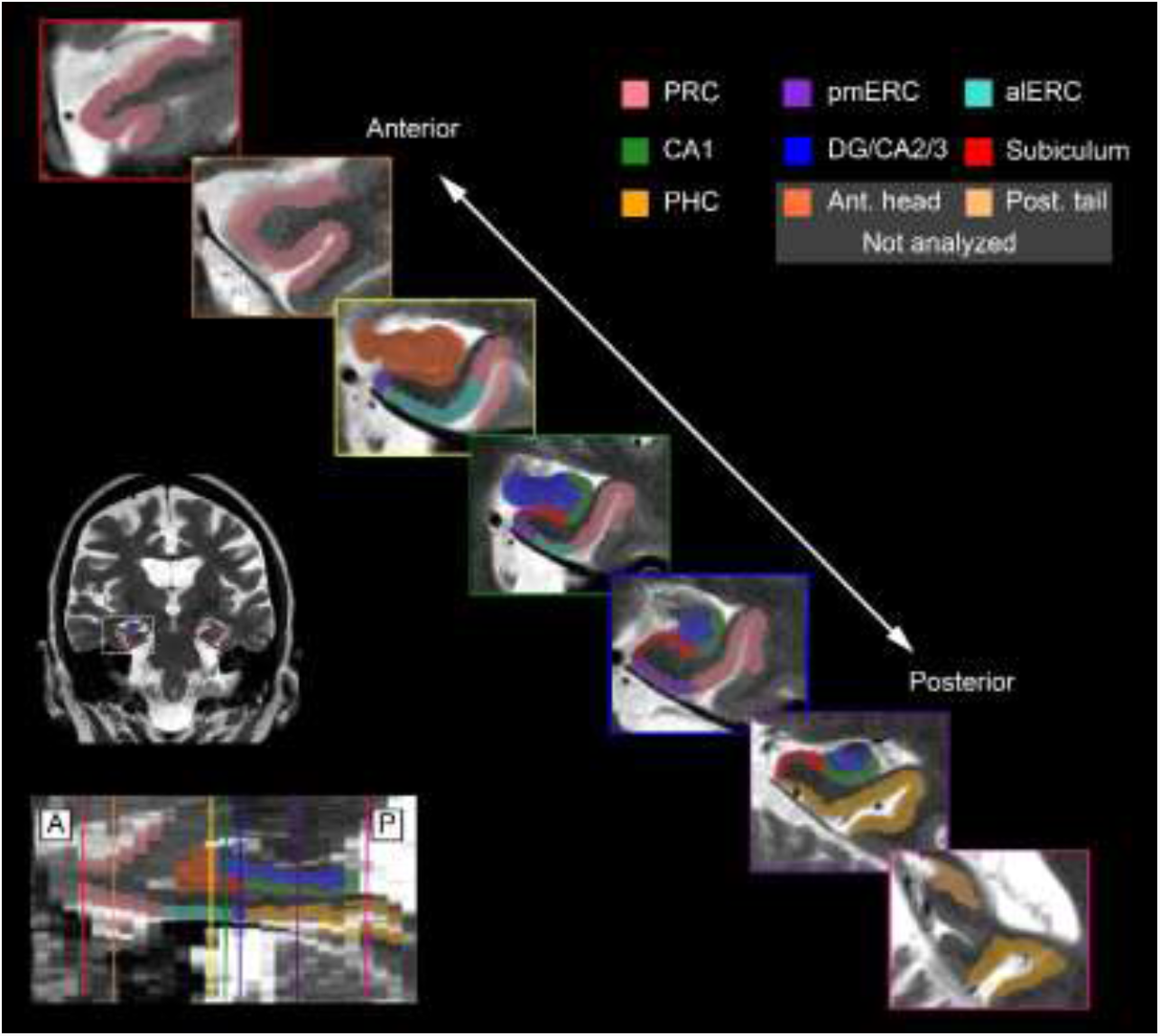
The modified version of the Olsen-Amaral-Palombo segmentation protocol used in the present study. Inset images depict coronal slices of the MTL taken at various points along the long axis of the hippocampus (as shown in the sagittal view in figure at bottom-left). Figure previously published in Olsen, Yeung et al. (2017), and reproduced with permission.

We considered these particular regions for two reasons. First, because these regions are directly connected to the alERC (Burwell, 2000; Suzuki & Amaral, 1994), we wished to explore if any observed alERC-behavior correlations were mediated by its inputs and outputs. Second, a number of these regions have been shown to be critically important in aspects of spatial memory, including object-location memory, and scene memory. PHC lesions have been shown to impair object-location memory (Bohbot et al., 1998; Malkova & Mishkin, 2003), and the PHC is reliably activated when viewing scenes (Epstein, Harris, Stanley, & Kanwisher, 1999). The hippocampus has long been known to have an important role in spatial representation (e.g. O’Keefe & Dostrovsky, 1971), and is theorized to support flexible representations of spatial/temporal arrangements of objects (Eichenbaum & Cohen, 2001) that underlie its role in scene memory and perception (Lee et al., 2005). The pmERC connects the PHC to the hippocampus, and direct recording work here suggests it is important for representing locations on a screen (Killian, Jutras, & Buffalo, 2012). The PRC is involved in combining object features into a conjunctive representation (Barense et al., 2012); here we are interested in how the alERC operates independently of the PRC. Our goal is to better understand how the alERC might contribute to all of these spatial/object memory processes that have long been associated with its surrounding MTL regions, and how volumetric differences in these regions in healthy older adults might affect those processes.

### Intra-and Inter-Rater Segmentation Reliability

Intra-rater reliability was established by comparing the segmentation of five randomly selected scans by the same rater (L.Y.) after a delay of 1-4 months. Inter-rater reliability was evaluated by comparing the segmentation of five randomly selected scans by a second rater (R.K.O) to those of L.Y. Both authors were blinded to MoCA score, task performance, and the identities of participants until after manual segmentation (including inter-and intra-rater reliability) was completed. Reliability was assessed using the intra-class correlation coefficient (ICC, which evaluates volume reliability) (Shrout & Fleiss, 1979) and the Dice metric (which also takes spatial overlap into account) (Dice, 1945), computed separately for each region in each hemisphere. ICC(3,k) was computed for intra-rater reliability (consistency) and ICC(2,k) was computed for inter-rater reliability (agreement). Dice was derived using the formula 2 * (area of intersecting region) / (area of original segmentation + area of repeat segmentation); a Dice overlap metric of 0 represents no overlap, whereas a metric of 1 represents perfect overlap. Intra-and inter-rater reliability results are shown in Table 2. These scores are comparable to reliability values reported in the literature for manual segmentation of hippocampal subfields and MTL cortices (Wisse et al., 2012; Yushkevich, Pluta, et al., 2015) and with our previous work (Olsen et al., 2013; Palombo et al., 2013).

**Table 2:** Inter-and intra-rater reliability measurements for manual segmentation. Dice was computed for both intra-and inter-rater agreement. ICC(3,k) was calculated for intra-rater and ICC(2,k) was computed for inter-rater reliability

| Subregion | Inter-rater: Dice |  | Inter-rater ICC |  | Intra-rater: Dice |  | Intra-rater: ICC |  |
| --- | --- | --- | --- | --- | --- | --- | --- | --- |
|  | Left | Right | Left | Right | Left | Right | Left | Right |
| CA1 | 0.88 | 0.87 | 0.94 | 0.95 | 0.74 | 0.66 | 0.92 | 0.91 |
| Subiculum | 0.85 | 0.84 | 0.89 | 0.88 | 0.67 | 0.66 | 0.81 | 0.85 |
| DG/CA23 | 0.91 | 0.90 | 0.94 | 0.99 | 0.75 | 0.73 | 0.91 | 0.96 |
| alERC | 0.86 | 0.85 | 0.96 | 0.86 | 0.72 | 0.73 | 0.87 | 0.71 |
| pmERC | 0.82 | 0.80 | 0.90 | 0.86 | 0.59 | 0.64 | 0.95 | 0.80 |
| PRC | 0.87 | 0.89 | 0.98 | 0.91 | 0.74 | 0.76 | 0.98 | 0.99 |
| PHC | 0.86 | 0.84 | 0.89 | 0.95 | 0.71 | 0.77 | 0.86 | 0.96 |

### Volume Correction for Head Size

All manually segmented region volumes were corrected for head size using a regression-based method, to account for differences in brain size between participants. Estimated total intracranial volume (eTIV) was derived from the whole-brain T1-weighted scans using FreeSurfer (v5.3) (Buckner et al., 2004). By regressing the volume of each region with eTIV, a regression slope β was obtained for each region (representing the effect of eTIV change on that region’s volume). Then, the volume of each region was adjusted by that participant’s eTIV using the formula Volume_adjusted_ = Volume_raw_ + β(eTIV_participant_ – eTIV_mean_). The head size correction was separately computed for each region in each hemisphere. In our previous work with this participant group (Olsen, Yeung et al., 2017), we did not observe an interaction between cognitive decline and hemisphere. Thus following this previous work, volumes were summed in each region across the two hemispheres, giving a single volume for each region for each participant.

### Eyetracker Setup

The experimental task was presented on a 21.2” monitor (36x30cm) at a resolution of 1024x768 pixels using Experiment Builder (SR Research, Mississauga, ON). Eyetracking measures were recorded using an EyeLink 1000 desktop-mounted eyetracker, sampling at a rate of 1000 Hz, with a spatial resolution of 0.01° and an accuracy of 0.25. Participants were positioned 55 cm away from the monitor, and placed their heads on a chin-rest to limit head motion. Nine-point calibration was performed prior to testing, and was repeated until the average gaze error was less than 1°, with no point having a gaze error exceeding 1.5°. Prior to each trial, a 1s drift correction was performed, with nine-point calibration being repeated if drift error exceeded 2°.

### Stimuli and Regions of Interest (ROIs)

Eight categories of computer-generated household scenes (bathrooms, bedrooms, backyard patios, dining rooms, garages, kitchens, living rooms, and office), were used in this study (Figure 2). All stimuli were created using Punch! home design software (Encore Software, Eden Prarie, MN). Each scene contained thematically appropriate objects, and all objects were unique to each individual scene. For each scene, we created two versions – a standard and an alternate. The alternate version of each scene was identical to the standard version in all respects, except for the location of a critical object within the scene. The standard version of the scene was used in all the “test” trials (i.e. it appeared in different test conditions for different participants as a result of counterbalancing). This arrangement allowed us to make direct comparisons between the same standard version of the scene, regardless of which test condition it appeared in for each individual participant. The design followed the counterbalancing procedures used in Ryan et al. (2000) and Smith et al. (2006). The standard version of each scene was also used in study trials whose scenes would be shown again in the repeated test condition. The alternate version of a scene was used for study trials whose corresponding test trial would be shown in the manipulated test condition. For instance, in Figure 2, the standard version of the scene is shown as the “test” scene in the manipulated test condition (center-bottom), whereas the alternate version of the scene is shown as the corresponding study trial above it. All of the scenes measured 1028x518 pixels, subtending the entire width and 2/3rds of the height of the display screen. The scenes were centered on the screen vertically.

**Figure 2:**
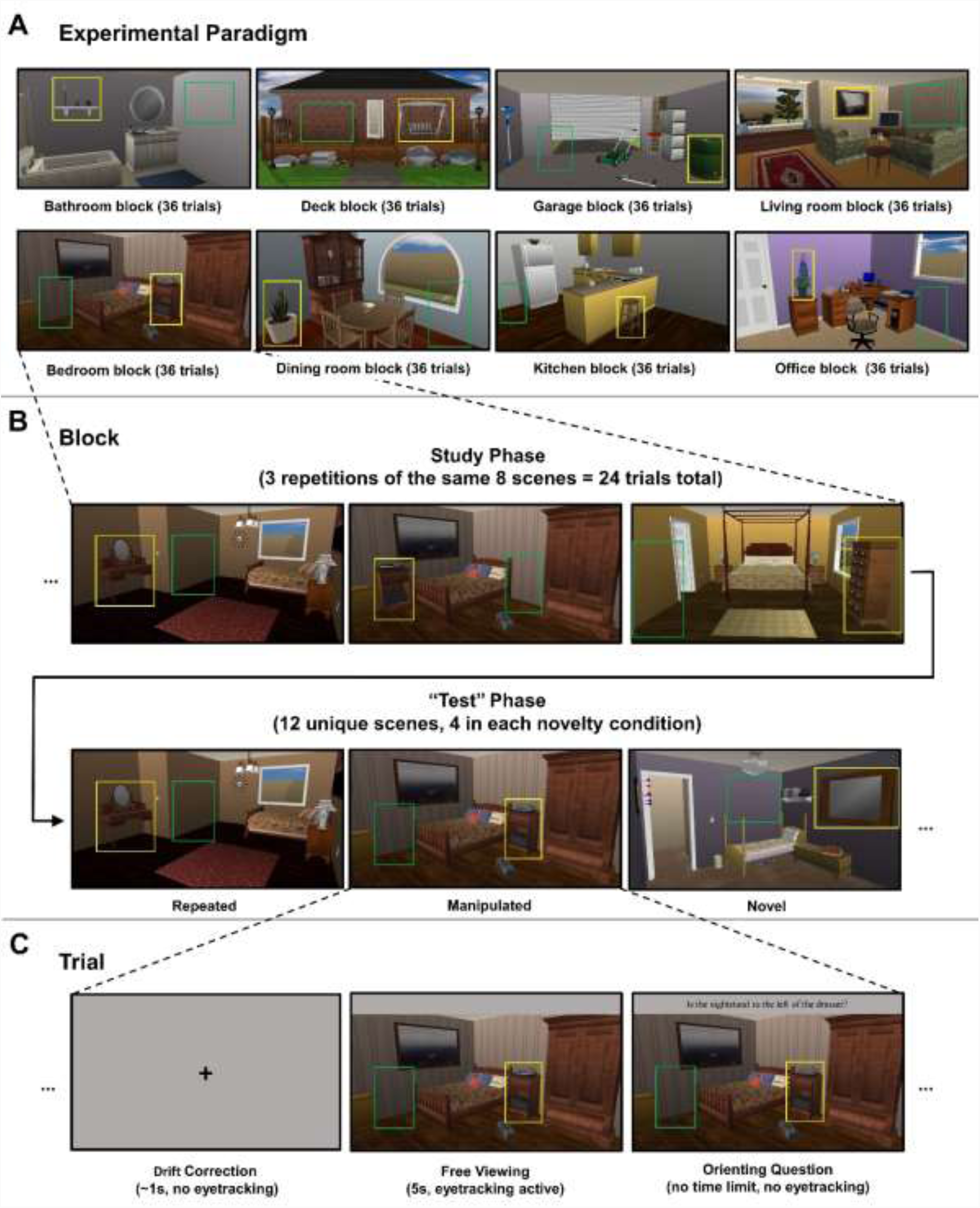
Schematic illustration of experimental paradigm. (A) Examples of each of the eight categories of scenes used in this study. (B) Arrangement of trials within each block. Each block had a study phase of 24 trials (8 unique scenes, repeated 3x), followed by a “test” phase of 12 trials (4 repeated scenes, 4 manipulated scenes, and 4 novel scenes). (C) Single trial timing. For all trials (in both study and “test” phases), after drift correction, participants freely viewed a scene for 5s, followed by a yes/no-orienting question directed to the critical object. For illustrative purposes, in all three panels, the critical object ROI (around the critical object) is shown in yellow, and the empty ROI (a similarly sized area covering the previous location of the critical object in manipulated scenes, or an empty location in repeated/novel scenes) is shown in green. Note that the ROIs were not visible to participants during the experiment.

Three rectangular regions of interest (ROI) were defined for each scene (Figure 3). The “whole scene ROI” (shown in red) encompassed the entire scene depicted and was uniformly 1028x518 pixels large. The “critical object ROI” (shown in yellow), was drawn to include only the critical object, and to minimize, as far as possible, the inclusion of parts of any other objects in the scene. The “empty ROI” (shown in green) covered an empty location in the scene, and was drawn to specifically minimize, as far as possible, parts of any other objects in the scene. Importantly, the empty ROI matched the location where the critical object ROI had been located during the study phase for *manipulated* scenes (which was simply an empty location on the scene for the *repeated* and *novel* scenes). Within each scene, the critical object ROI and empty ROI were similarly, but not identically, sized. This was necessary to ensure that the ROIs did not include, as far as possible, any part of any other objects in the scene, which might receive additional fixations during the viewing period that are not directed to the critical object, or the empty location. Note that no comparisons were made between the critical object ROI and the empty ROI; rather all comparisons were within the same ROI across conditions. Across the entire stimulus set, the mean critical object ROI had an area of 36,122 pixels (6.80% of the scene, SD = 20,853 pixels), whereas the mean empty ROI had an area of 37,575 pixels (7.08% of the scene, SD = 17,778 pixels).

**Figure 3:**
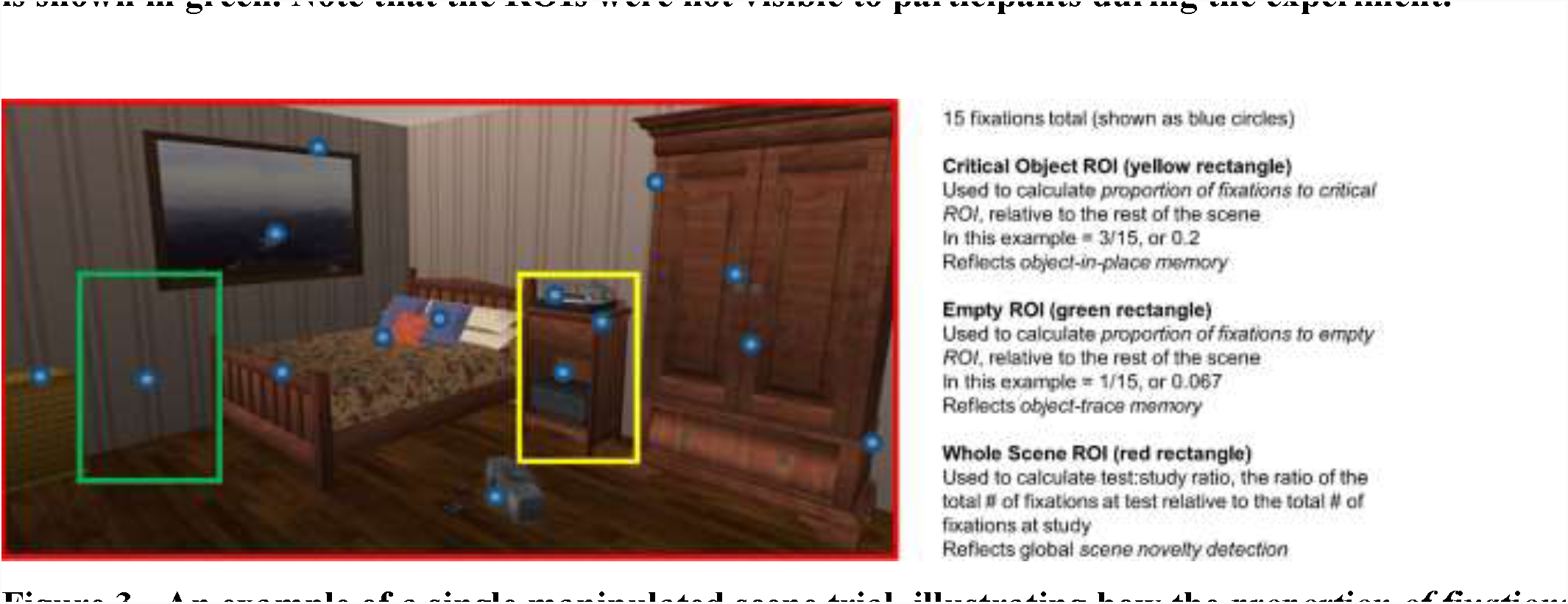
An example of a single manipulated scene trial, illustrating how the *proportion of fixations* outcome variables were calculated. Note that the ROIs and fixations were not visible to participants as they performed the task.

### Eyetracking Task

We employed an eyetracking-based paradigm assessing processing of objects within scenes, as assessed at varying levels of novelty, and examined how volumetric differences in alERC (and other MTL/hippocampal regions) affected object-in-place memory and object-trace memory (Figures 2 and 3). In each trial, participants incidentally viewed computer-generated scenes, depicting household locations (e.g. bedrooms, kitchens, etc.), for 5s (Figure 2C). After viewing each scene, participants were asked to respond to a yes/no orienting question (appearing above the scene), directing attention to a critical object in the scene (e.g. “Is the nightstand to the left of the dresser?”). This followed the examples of Ryan et al., (2000) and Hannula, Tranel, & Cohen (2006) who also used orienting questions to direct viewers’ attention to a critical object within a scene. No time limit was imposed on answering this orienting question. Visual fixations made within the three ROIs (Figure 3) were recorded during the 5s viewing period, but not during the subsequent period when participants were asked to respond to the orienting question. A brief eyetracker drift correction (<1s) was performed between each trial.

The experiment was organized into eight blocks of 36 trials each; all the scenes in each block depicted the same type of location (e.g. one block consisted entirely of bedrooms, another block entirely of kitchens, etc.) (Figure 2A). Each block included a study phase of 24 trials (eight unique scenes viewed three times each, all eight scenes were viewed at least once before any scenes repeated), followed by a “test” phase of 12 trials (Figure 2B). It is important to note that participants were not informed of any distinction between the study and “test” phases, as the task instructions were the same across all trials. In each “test” phase, there were four scenes in each of three test conditions (i.e. 12 trials total in each “test” phase), which differed in the degree of novelty (Figure 2B). The three test conditions were: 1) *repeated scenes*, which were identical versions of the scenes presented during the study phase, 2) *manipulated scenes*, which were identical to a scene presented during the study phase, except that the critical object had moved to a different location in the scene, and 3) *novel scenes*, which were not seen during the study phase, but depicted the same type of scene as the rest of the block (e.g. a kitchen in a block of kitchens).

Each of the repeated or manipulated scenes shown during the “test” phase corresponded to one specific scene viewed three times during the study phase. For example, in Figure 2B, the repeated scene (bottom left) is identical to a scene viewed during the study phase (top left), whereas the manipulated scene (bottom center) is the same as the studied scene (top center), with the exception that the critical object (the nightstand) was moved from the middle of the room to the left side of the room. Further, the orienting question for each specific scene (including both versions of the scenes used for the manipulated test condition) remained the same across all repetitions, as did the correct answer to that orienting question. For instance, in Figure 2, the orienting question for the middle scene (with the nightstand) was “Is the nightstand to the left of the dresser?” The answer to this question was the same (“yes”) for both the alternate version of the scene shown during the study phase and the standard version of the scene shown during the “test” phase as a manipulated scene.

Across participants, the condition in which a particular scene appeared (see “Stimuli and Regions of Interest (ROIs)” section above), the ordering of the blocks, and the correct response to the orienting question for each scene, were all counterbalanced. For instance, in the nightstand scene we have been using as an example, half of the participants received the alternate orienting question of “Is there a stereo on top of the nightstand?” to which the correct answer is “no” in both versions of the scene.

### Eyetracking Outcome Variables

We defined three eyetracking-based outcome variables for each test condition, based on similar measures previously employed in other studies of object-scene memory (Ryan et al., 2000, 2007, Ryan & Cohen, 2004b, 2004a; Smith et al., 2006; Smith & Squire, 2008) (Figure 3). Our first primary outcome variable was the *proportion of fixations to the critical object ROI*. We used this measure of viewing to assess object-in-place memory. This measure was calculated for each individual trial by dividing the number of fixations made to the critical object ROI by the total number of fixations made to the whole scene ROI, and then by averaging over all the trials in a single test condition (i.e., repeated, manipulated, or novel) for each participant. In the novel scene condition, participants would not have known which object was the critical object (as they had yet to be shown the orienting question matching that scene). However, in the repeated and manipulated scene conditions, participants would have previously studied an identical (or nearly-identical) version of the scene, and thus, would have had the opportunity to associate the critical object mentioned in the orienting question with its spatial location in that scene. We were particularly interested in the how the proportion of fixations directed to the critical object ROI differed across the three test conditions. The difference in viewing the critical object ROI between the repeated/manipulated conditions and the novel condition reflects memory for having previously viewed the critical object. Further, the difference in viewing the critical object ROI between the repeated and manipulated conditions reflects memory for the location of the critical object within that scene (i.e. object-in-place memory). We hypothesized that alERC and PHC volume would be related to this measure, based on their role in object-location memory. Based on its role in spatial memory, hippocampal subfield volumes may be related to this measure too.

Our second outcome variable was the *proportion of fixations to the empty ROI* and was used to assess object-trace memory. This was calculated in a similar manner as the average proportion of fixations to the critical object ROI. For each trial, the number of fixations to the empty ROI was divided by the number of fixations to the entire scene. Then, we averaged this proportion over all the trials in each test condition separately (Figure 3). The difference in viewing between the manipulated condition, and the repeated/novel conditions provided a measure of object-trace memory: increased viewing to this otherwise-empty location reflects memory for the previous location of the critical object. By the same rationale as above, we also hypothesized that alERC, PHC, and hippocampal subfield volumes may be related to this measure.

Our third outcome variable was the *test:study ratio*. This was used to assess global scene novelty detection, as well as to see if the differences in viewing to the critical object we observed could be explained by the broader relationship between viewing the scene as a whole and MTL regional volumes. This measure was calculated by taking the average number of fixations made to the whole scene ROIs for all the scenes in a given test condition (repeated, manipulated, or novel) and normalizing by the average number of fixations made to all the scenes during their initial presentation in the study phase. The number of fixations made to scenes during the first presentation in the study phase within each block (when all the scenes were entirely novel) served as a baseline for how many fixations a particular participant would make to an entirely novel scene. This followed the procedure we employed in our previous work (Yeung, Ryan, Cowell, & Barense, 2013) to derive a normalized eyetracking-based measure of novelty that controlled for absolute differences in the number of fixations between participants. Because more fixations are made to novel stimuli compared to previously-viewed stimuli (Althoff & Cohen, 1999), this variable served as a measure of novelty detection for the scene as a whole (i.e. scene memory). A score of 1 or more here indicates that scenes in a given test condition were being treated as novel (i.e. the same number of fixations were made as compared to when the scene was entirely novel during the first presentation of the study phase). In contrast, a score of less than 1 indicates that the scenes were visually sampled as though they had been previously seen (i.e. fewer fixations were made as compared to when the scene was entirely novel). In contrast to the previous two outcome measures, we did not predict that any MTL brain regions would be related to this measure. Although the PHC and hippocampus are implicated in spatial processing, the repeated presentation of identical scenes would allow lower-level visual areas to assess global scene novelty, without the involvement of the PHC or hippocampus (Cowell et al., 2010). This is consistent with previous work demonstrating that amnesiac individuals with MTL damage do not show any impairments in the eye-movement based memory effect for scenes (Ryan et al., 2000).

### Statistical Analysis

Repeated-measures ANOVAs and planned paired-samples t-tests were used to identify differences in each of the three behavioral outcome variables (proportion of fixations to the critical object ROI, proportion of fixations to the empty ROI, test:study ratio; Figure 3), in each test condition (i.e. novel, manipulated, and repeated). To assess the importance of each brain region on the primary behavioral outcome variable, multiple regression was employed with each outcome variable as the dependent variable and the volumes of the seven brain regions as predictors, for each test condition separately.

For test conditions where brain region volumes were significant predictors for a behavioral outcome variable, we were interested in whether that behavior predicted variation in brain region volume above the effects captured by existing measures of cognitive decline (e.g. the MoCA) or aging. That is, we asked: Does our outcome measure have predictive value beyond existing measures? To this end, additional multiple regression analyses were conducted with the dependent variable, MoCA and age as predictors for the volume of those brain regions. All statistical tests were two-tailed, and conducted at α=0.05. Mauchly’s test of sphericity was applied to repeated-measures ANOVAs; when the assumption of sphericity was violated, the Greenhouse-Geisser correction was applied. Multiple regressions were tested for multicollinearity; residual plots were inspected to check for non-linearity and heteroscedasticity.

## Results

### Visual Sampling and Behavioral Responses

#### Orienting Question

We first looked at whether participants differed in their ability to assess the spatial relations among objects in the scene (i.e. whether they were able to accurately answer the orienting question). A repeated-measures ANOVA showed no effect of test condition (repeated, manipulated, novel) on the accuracy of responses to the orienting question, F(2,58) = 0.581, p = 0.563, η^2^ = 0.020. Indeed, accuracy to the orienting question was uniformly high across all three test conditions: mean = 92.0%, SD = 8.1% for the repeated condition, mean = 90.5%, SD = 7.2% for the manipulated condition, and mean = 91.8%, SD = 7.0% for the novel condition. That is, across all test conditions, participants did not differ in their ability to perceptually assess spatial relations among presented objects in a scene.

#### Object-in-Place Memory

To investigate object-in-place memory, we ran a repeated-measures ANOVA investigating the effect of test condition (repeated, manipulated, novel) for the proportion of fixations directed to the critical object ROI. These showed a main effect of test condition for the proportion of fixations directed to the critical object ROI, F(2,58) = 13.874, p < 0.001, η^2^ = 0.324 (Figure 4A). Next, we compared the proportion of fixations to the critical object ROI in the repeated/manipulated scenes (where the critical object was known due to previous viewings of the scene during the study phase), to the same measure in the novel scenes (where the critical object is unknown as that scene was not shown in the study phase). Paired-samples t-tests showed that a greater proportion of fixations were directed to the critical object ROI for repeated scenes than novel scenes, t(29) = 2.342, p = 0.026, d = 0.453, and for manipulated scenes than novel scenes, t(29) = 6.301, p < 0.001, d = 1.009. This suggests a memory effect for having previously viewed the critical object in the repeated and manipulated conditions. We further compared the proportion of fixations to the critical object ROI between the repeated and manipulated conditions, to explore whether there was memory for the location of the critical object within the same scene. We found that the difference in the proportion of fixations to the critical object ROI between the repeated and manipulated conditions was significant, t(29) = 2.583, p = 0.015, d = 0.431 suggesting memory for the location of the critical object within the scene, beyond simply memory for the critical object (Figure 4A).

**Figure 4:**
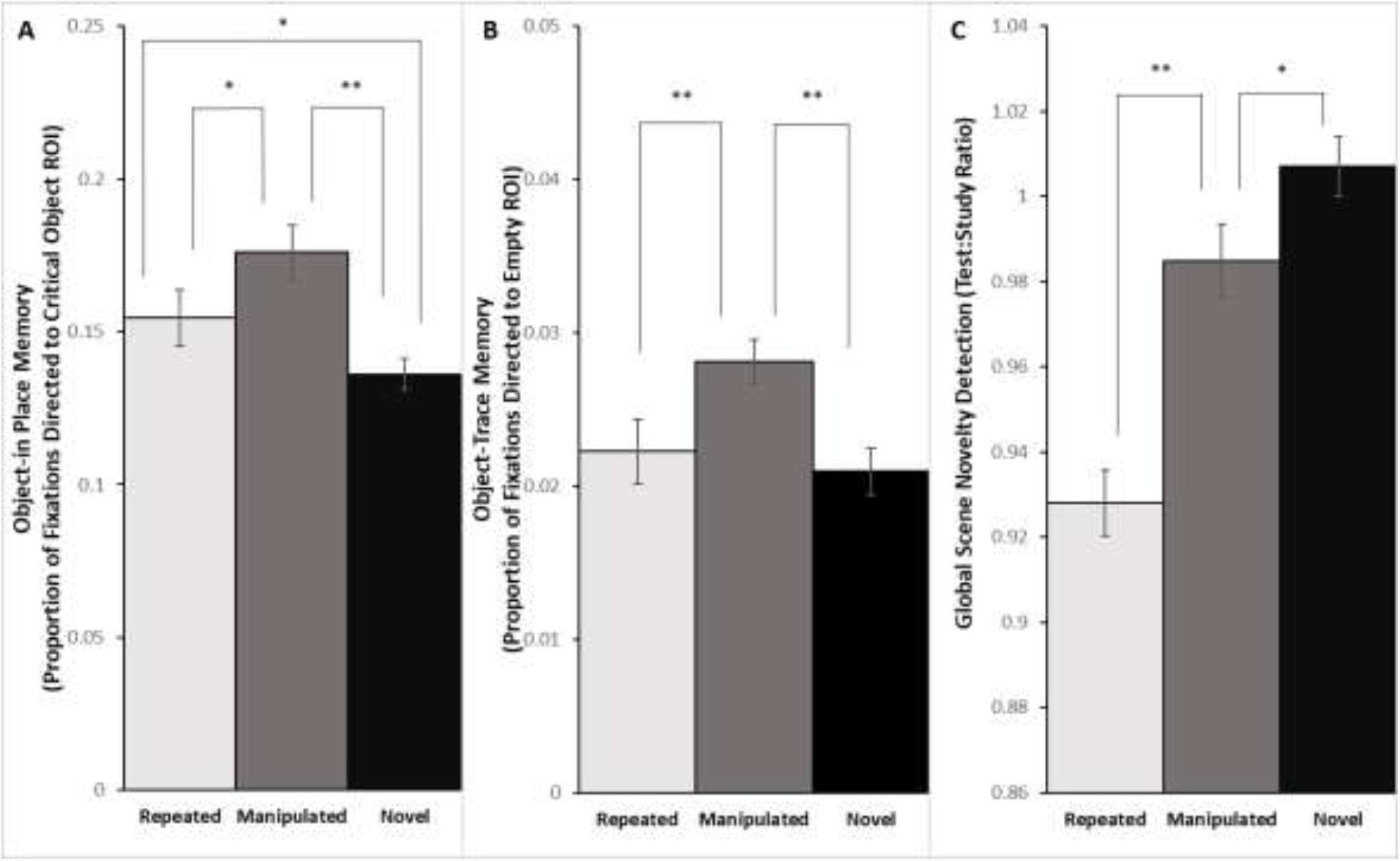
Behavioral eyetracking results. (A) The proportion of fixations directed to the critical object ROI relative to the entire scene, a measure of object-in-place memory. (B) The proportion of fixations directed to the empty ROI relative to the entire scene, a measure of object-trace memory. (C) The test:study ratio, a measure of global scene novelty detection. ** p < 0.01, * p < 0.05. Error bars represent SEM. N=30 for all conditions.

#### Object-Trace Memory

To investigate object-trace memory, we ran a repeated-measures ANOVA investigating the effect of test condition (repeated, manipulated, novel) for the proportion of fixations directed to the empty ROI. These showed a main effect of test condition F(2,58) = 6.410, p = 0.003, η^2^ = 0.181 (Figure 4B). We evaluated object-trace memory across different test conditions by comparing the proportion of fixations to the empty ROI in the manipulated scenes (where the critical object had been located during the study phase), to the repeated/novel scenes (where it was only an empty location in the scene). Paired-samples t-tests showed that a greater proportion of fixations were directed to the empty ROI in the manipulated condition compared to either the repeated condition t(29) = 2.912, p = 0.007, d = 0.592, or the novel condition, t(29) = 3.760, p < 0.001, d = 0.881. In contrast, there was no difference between the repeated and novel conditions, t(29) = 0.535, p = 0.597, d = 0.128 (Figure 4B). The greater degree of viewing to the empty ROI in the manipulated scenes suggests the presence of object-trace memory for the location where the critical object was previously located.

#### Global Scene Novelty Detection

To investigate global scene novelty detection, we ran a third repeated-measures ANOVA investigating the effect of test condition (repeated, manipulated, novel) on the test:study ratio. These showed a main effect of test condition for the test:study ratio (Figure 4C), F(2,58) = 36.786, p < 0.001, η^2^ = 0.559. Paired-samples t-tests showed that the test:study ratio was significantly higher for manipulated scenes compared to repeated scenes, t(29) = 5.649, p < 0.001, d = 1.263 and in turn, significantly higher for novel scenes compared to manipulated scenes, t(29) = 2.537, p = 0.017, d = 0.517 (Figure 4C). These results indicate a larger eye-movement based memory effect for the repeated scenes (as they received fewer fixations than the novel condition), and a smaller effect for the manipulated scenes. This replicates the eye-movement based memory effect for scenes (e.g. Ryan et al., 2000), as well as for objects (Yeung et al., 2017, 2013) as observed in our previous work.

### Relationship between Viewing Measures and MTL Regional Volumes

#### Object-in-Place Memory

Next, we examined the influence of differences in MTL/hippocampal subregion volumes on viewing to the critical object ROI for each condition, our measure of object-in-place memory. Multiple regression analysis using the seven brain region volumes as predictors (Table 5A) revealed that only alERC and PHC volumes were significant predictors for the proportion of fixations made to the critical object ROI, and only for the manipulated condition (alERC: t(29) = 2.61, p = 0.02, β = 0.46, sr = 0.41; PHC: t(29) = 2.50, p = 0.02, β = 0.48, sr = 0.39, statistical tests for all predictors shown in Table 5A). That is, greater alERC and PHC volume predicted a greater proportion of fixations to the critical object, but only when the critical object was moved from its position in a previously studied scene (Figure 5 illustrates this relationship graphically). This suggests that the volumes of these two regions were related to the strength of the memory for the critical object and its spatial location within the scene.

**Table 2:**
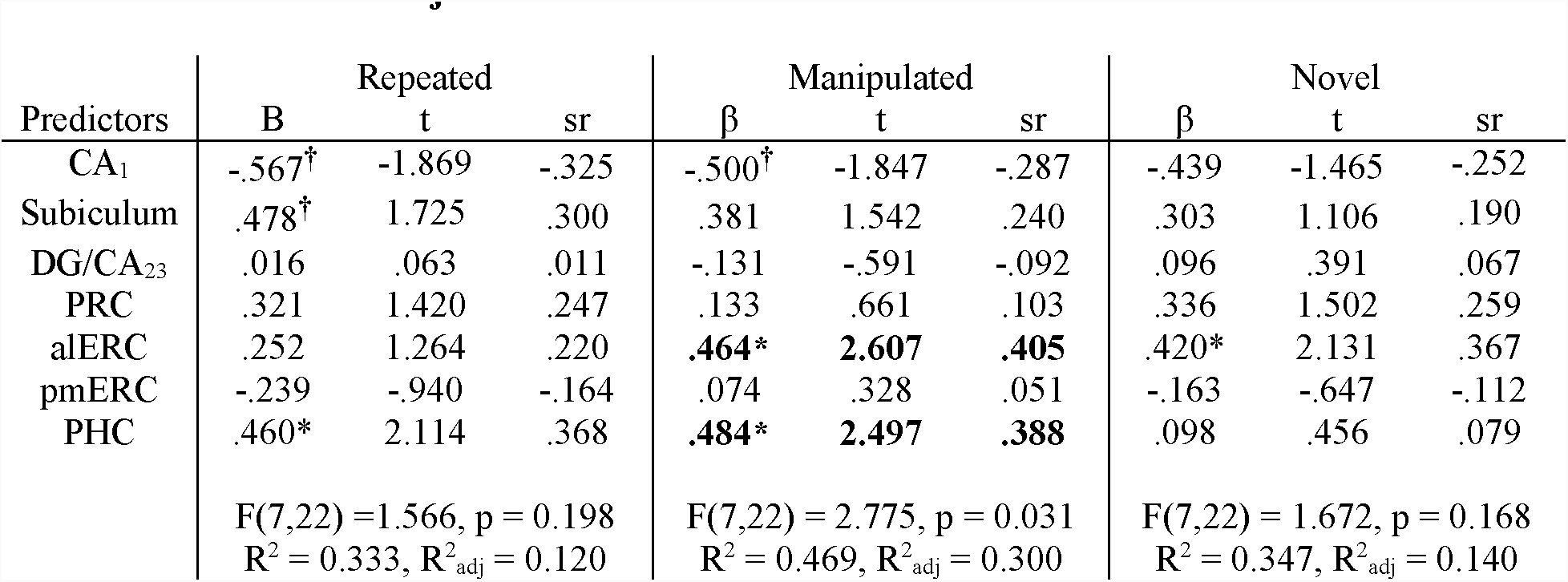
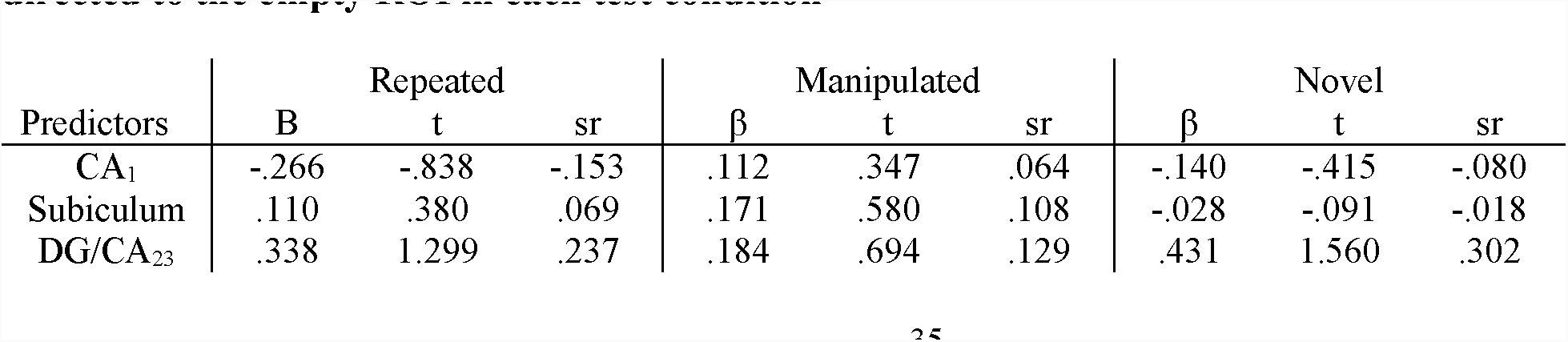
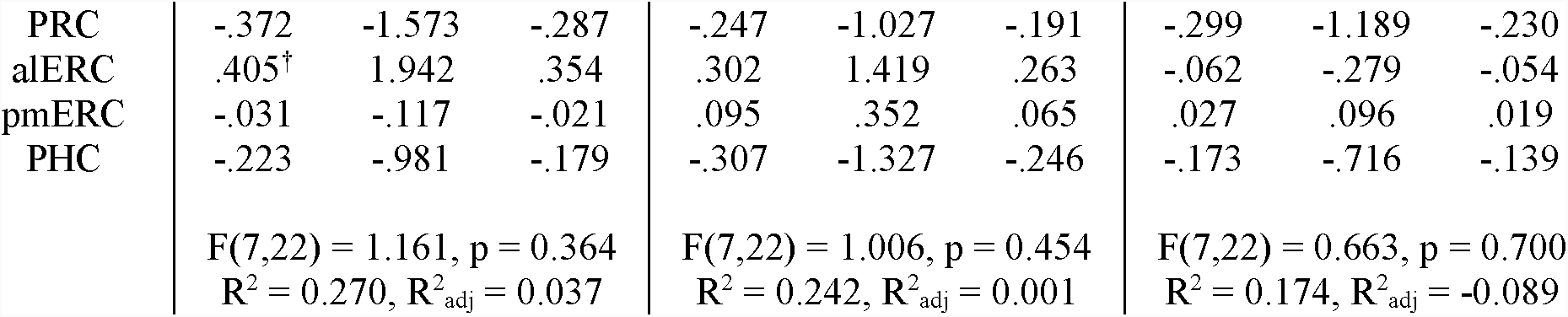
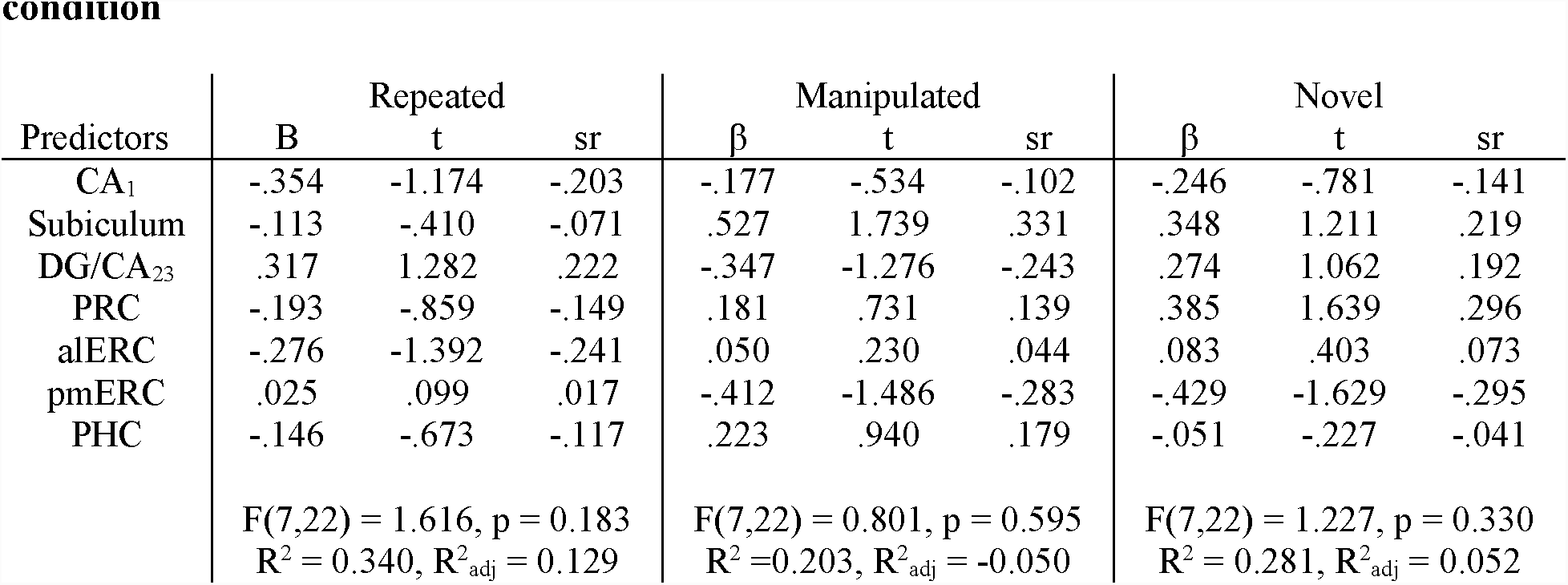
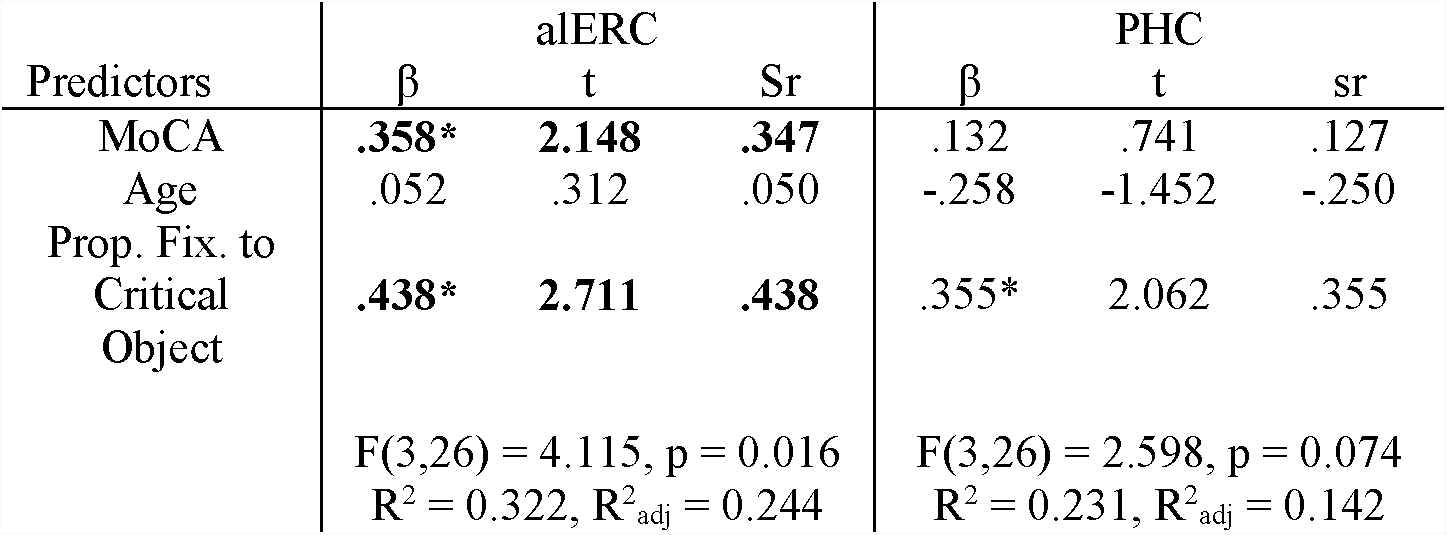
Summary tables for multiple regression analyses. In all three tables, multiple regression models were run separately for trials in each test condition. Each model is shown in its own column. † p < 0.1, * p < 0.05, ** p < 0.01 (reflect-tests for significance of each predictor) **(A) Multiple regression analyses with brain region volumes as predictors for the proportion of fixations directed to the critical object ROI in each test condition** **(B) Multiple regression analyses with brain region volumes as predictors for the proportion of fixations directed to the empty ROI in each test condition** **(C) Multiple regression analyses with brain region volumes as predictors for the test:study ratio in each test condition** **(D) Multiple regression analyses with MoCA, age and viewing to the critical object ROI in the manipulated condition (object-in-place memory) as predictors for alERC and PHC volume**

**Figure 5:**
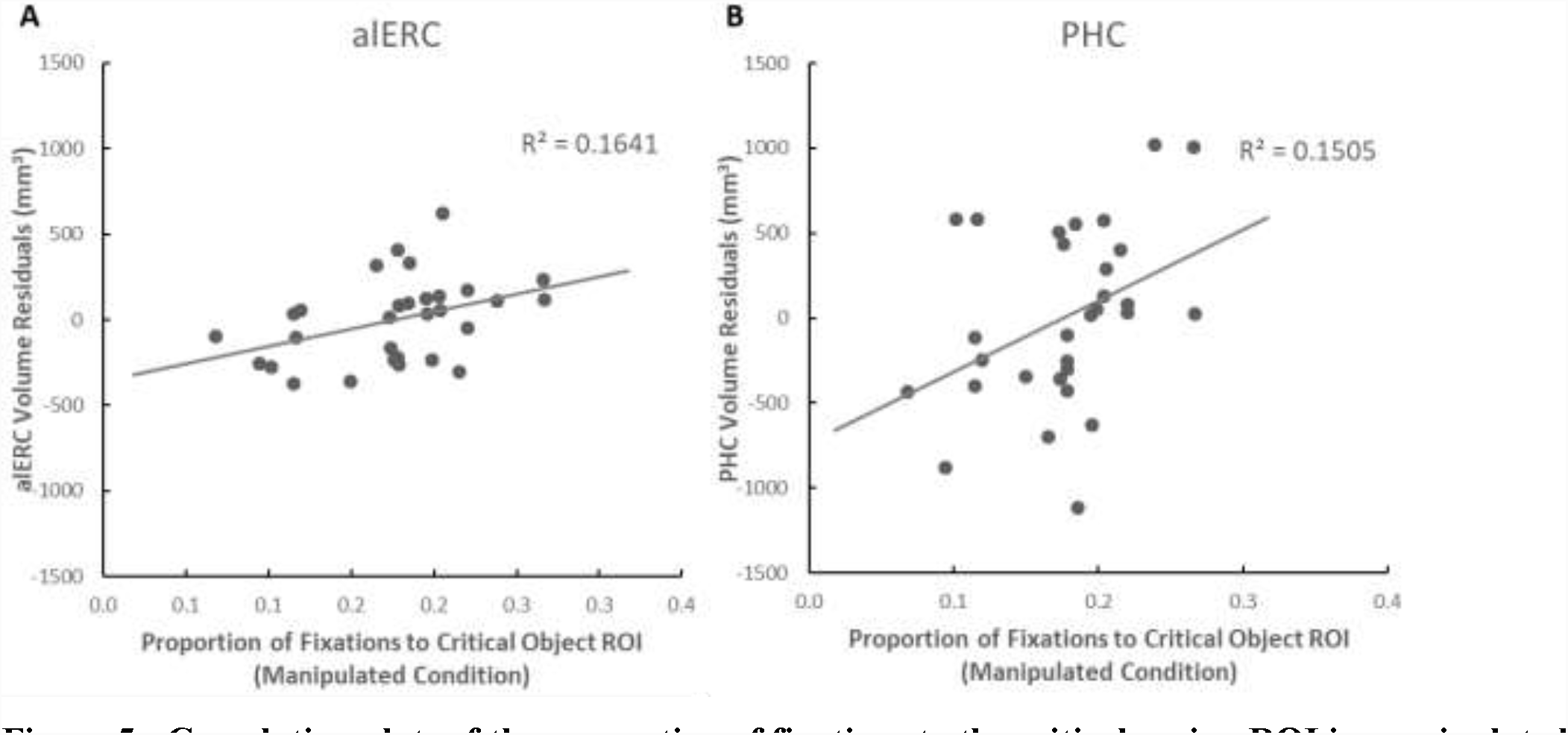
Correlation plots of the proportion of fixations to the critical region ROI in manipulated scenes relative to the (A) alERC residuals and (B) PHC residuals. These depict solely the contribution of the alERC and PHC predictors in Table 5A (i.e. with the contribution of other regions removed), demonstrating a correlation of alERC and PHC structural volumes with an eye-movement based measure of object-in-place processing.

#### Object-Trace Memory

Next, we investigated the effect of our MTL regional volumes on viewing to the empty ROI for each condition, our measure of object-trace memory. None of the multiple regression models predicting the proportion of fixations to the empty ROI were significant for any condition (Table 5B). That is to say, MTL regional volume differences did not seem to relate to object-trace memory.

#### Global Scene Novelty Detection

We also calculated the relationship between MTL regional volumes and global scene novelty detection, as measured by the test:study ratio. Consistent with our previous findings (Yeung et al., 2017), none of the tested regions significantly predicted the test:study ratio for any condition (Table 5C); i.e. MTL volume differences did not affect eye movement based measures of novelty for the scene as a whole. Note that this is consistent with previous reports showing that for amnesiacs, eye movement repetition effects were normal for faces (Althoff et al., 1998; Olsen et al., 2016) and scenes (Ryan et al., 2000)when procedures are akin to those used here (e.g., multiple repetitions, relatively extended viewing time per stimulus presentation.

#### Predictive Effects of Viewing Compared to MoCA and Age

Having established a connection between alERC/PHC volume and viewing behavior towards the critical object ROI in the manipulated condition, we next asked whether viewing behavior predicted change in the volumes of those regions, above and beyond the effects of MoCA or age (Table 5D). This question is important to ascertain whether the viewing behavior we report might be indicative of alERC/PHC volume differences not accounted for by existing cognitive measures. Even with MoCA and age as predictors in the same model (i.e. having accounted for the variance in alERC and PHC volume they explain), the proportion of fixations to the critical object ROI in the manipulated condition was a significant predictor for alERC volume (t(29) = 2.71, p = 0.01, β = 0.44, sr = 0.44). That is to say, the proportion of fixations to the critical object ROI for manipulated scenes explained a significant variance in alERC volume even after having accounted for the effects of cognitive decline (as assessed by the MoCA). In contrast, the multiple regression model predicting PHC volume using the same predictors only trended towards significance, F(3,26) = 2.60, p = 0.074, thus caution is to be taken in drawing strong conclusions here regarding the effects of individual predictors.

## Discussion

The alERC is an important region in the early progression of dementia: it is coterminous with regions where AD-related tau pathology first appears in the MTL (Braak & Braak, 1991; Khan et al., 2014), and its volume is reduced in individuals who score below threshold on a neuropsychological test sensitive to AD-related cognitive decline (Olsen, Yeung et al., 2017). Despite the clinical importance of the alERC, its cognitive functions remain poorly explored. In this study, we investigated how alERC volume differences in older adults related to visual processing of objects in scenes. We found that alERC volume predicted object-in-place memory (the association of an object with a particular location in a scene). In particular, alERC volume tracked viewing to a critical object associated with a scene only when that object had moved to a new location. This is the first human study to suggest that the alERC plays a role in supporting the spatial associations of an object within a scene. It also demonstrates a remarkably specific link between brain structure and cognitive behavior: differences in alERC volume are connected to how people direct their gaze when viewing a scene.

Few cognitive models differentiate between the subregions of the human entorhinal cortex. One model that makes this distinction - the PMAT model - proposes that the alERC (combined with the PRC from where it receives most of its inputs) is part of a system that represents the properties of unique entities (i.e. objects). In contrast, the pmERC belongs to a different system representing the spatial, temporal or causal relationships between entities (Ritchey et al., 2015). Previous human neuroimaging studies support this model, showing that lateral regions of the ERC are more active when differentiating between objects that differed in their features versus objects that differed by location (Reagh et al., 2018; Reagh & Yassa, 2014), and when differentiating between faces/objects compared to scenes (Berron et al., 2018; Schultz et al., 2012). This distinction is also supported by some rodent studies showing object-responsive cells in the homologous LEC (Deshmukh, Johnson, & Knierim, 2012; Deshmukh & Knierim, 2011), and impaired object novelty detection in LEC-lesioned rats (Hunsaker, Chen, Tran, & Kesner, 2013). However, some argue that the LEC may also represent the spatial location of objects (Connor & Knierim, 2017; Knierim, Neunuebel, & Deshmukh, 2013), based on studies that show rodent LEC lesions impair spatial navigation (Kuruvilla & Ainge, 2017; Van Cauter et al., 2013) and object-context memory (Wilson et al., 2013). Direct recording from rodent LEC show neurons with place fields for locations that previously held objects (Deshmukh & Knierim, 2011; Tsao et al., 2013), and LEC neurons that are responsive to object and spatial dimensions (Keene et al., 2016). Our results provide the first evidence that the human alERC (like the rodent LEC) also plays a role in supporting the spatial relations of an object within a scene, suggesting it may have a larger role than attributed to it by the PMAT model.

These data are consistent with the notion that there is a representational hierarchy throughout the ventral visual stream that extends into the MTL (Cowell et al., 2010; Saksida & Bussey, 2010). Moving anteriorly through the ventral visual stream, successive regions support a hierarchy of stimulus representation, such that an object’s low-level features are represented in early posterior regions, whereas increasingly complex conjunctions of object features are represented in more anterior regions (Desimone & Ungerleider, 1989; Riesenhuber & Poggio, 1999; Tanaka, 1996). Recent evidence suggests that the PRC, which receive inputs from the anterior regions of the ventral visual stream, contains representations of objects as a whole (Barense et al., 2012; Bussey, Saksida, & Murray, 2002; Erez, Cusack, Kendall, & Barense, 2016). This study’s results suggest that alERC (receiving inputs from the PRC) supports even more complex object representations integrating spatial information about how an object relates to its environment. In turn, these alERC representations can be further combined into hippocampal-dependent representations of scenes comprised of flexible relations among objects (see Lee, Yeung, & Barense, 2012 and Olsen, Moses, Riggs, & Ryan, 2012, for review).

How might we reconcile our results with reports that the human lateral ERC is not involved in spatial processing? For instance, Reagh & Yassa (2014) reported only minimal BOLD activity in human lateral ERC when trying to ascertain whether an identical object had moved slightly in a blank field. In contrast, we report that human alERC volume affects how much we fixate on an object that has moved within a particular scene. These contrasting results suggest that the alERC’s role may not be directly representing the location of an object, but rather lies in representing the spatial relations of an object within a scene (or relative to other objects). Combined with our previous work showing that alERC volume predicted viewing to configurally-important regions of objects (i.e. processing the spatial relations between parts of an object; Yeung et al., 2017), these results suggest the human alERC is involved in representing some spatial properties of objects. Speculatively, the alERC may be a region where distinct information from the two systems of the PMAT system begin to converge (Suzuki & Amaral, 1994).

There is a fascinating body of work looking at direct recording from the monkey ERC during free-viewing of visual images, which reported cells responsive to visual exploration (analogous to cells in the rodent ERC that respond selectively to ambulatory exploration). For example, there are reports of monkey ERC visual grid cells, which fire in response to visual fixations to a series of regions arranged in a hexagonal pattern (Killian et al., 2012; Meister & Buffalo, 2018). These cells are analogous to rodent ERC grid cells that fire when a rat traverses a set of regions arranged in a similar hexagonal pattern (Hafting, Fyhn, Molden, Moser, & Moser, 2005). Direct recording studies have also discovered monkey ERC “saccade direction cells”, which fire in response to saccades moving in specific directions (Killian, Potter, & Buffalo, 2015), analogous to rodent ERC head direction cells (Sargolini et al., 2006). The correspondence of these studies suggests that visual exploration in monkeys (and by extension, humans as well) is analogous to physical exploration in rodents (Ellard, 1998; Whishaw, 2003) in its neural representation. Our work builds on the literature in two important ways. Firstly, we demonstrate a unique correspondence between visual exploration behavior and ERC volume, suggesting there may be population-level effects among ERC neurons on visual behavior (see also Meister & Buffalo, 2018). Secondly, we are the first group (to our knowledge) to look at the connection between the ERC and viewing behavior in humans.

We also found that PHC volume was significantly related to viewing directed to the critical object in manipulated scenes. This is consistent with reports from the lesion (e.g. Bohbot et al., 1998; Malkova & Mishkin, 2003) and functional imaging literature (e.g. Buffalo, Bellgowan, & Martin, 2006; Cansino, Maquet, Dolan, & Rugg, 2002; Düzel et al., 2003; Maguire, Frith, Burgess, Donnett, & O’Keefe, 1998) showing PHC involvement in object-location memory. The PHC has also shown to be activated in response to navigationally-relevant (Janzen & Van Turennout, 2004) or spatially-defining objects (Mullally & Maguire, 2011); our results may reflect the influence of the PHC on viewing behavior to the critical object in relation to other scene elements. Our data also suggest this PHC function may involve the alERC, potentially reflecting the combination of information from both networks of the PMAT model. In contrast, we did not observe any effect of pmERC volume on viewing behavior. This was surprising to us, given reports of monkey visual grid cells in posterior ERC (Killian et al., 2012). However, it is theorized that the rodent medial ERC is more involved in spatial navigation (Knierim et al., 2013), and our task lacks navigational demands. It is also possible that as different grid cells code for location at different scales and orientations (Killian et al., 2012), there is an intrinsic redundancy that may help the pmERC resist behavioral changes as the result of slight volume loss.

Numerous experiments have shown that hippocampal function is related to ongoing visual exploration (e.g. Voss, Bridge, Cohen, & Walker, 2017, see Hannula, Ryan, & Warren, 2017 for review). Whereas healthy participants typically showed a relational manipulation effect of increased viewing to manipulated regions of scenes (as we also observed in this study), amnesic cases who presented with lesions to the hippocampus (and/or surrounding MTL) did not (Ryan et al., 2000; Ryan & Cohen, 2004a). Moreover, amnesic cases with hippocampal damage did not show increased viewing towards faces associated with a particular scene (Hannula, Ryan, Tranel, & Cohen, 2007), and in fact demonstrated an altered pattern of visual exploration of faces (Olsen et al., 2015). Neuroimaging studies have also shown hippocampal BOLD activity related to eyetracking-based measures of memory. Hippocampal BOLD activity was correlated with increased fixations to faces associated with a particular scene (Hannula & Ranganath, 2009), and fixations directed to similar locations in structurally-similar scenes (Ryals, Wang, Polnaszek, & Voss, 2015); in both cases, hippocampal activation was observed even when participants were unable to explicitly articulate the similarities they observed. Given these results, we expected that hippocampal subfield volumes would relate to individual differences in viewing towards the critical object in our study; however, we did not observe such a relationship in our data. This result might be accounted for by differences in methodology and population: our study looked at the relationship between cognitive performance and regional brain volumes in relatively healthy older adults, whereas previous work looked at fMRI activity in younger adults, or cognitive performance in amnesic cases with specific lesions. There are age-related differences in eyetracking measures; for instance, increased viewing to the critical region of manipulated scenes has been consistently reported in younger adults (Ryan et al., 2000, 2007; Smith et al., 2006), whereas this is not the case for older adults (Ryan et al., 2007). Further, brain lesions can lead to greater damage in white matter tracts between MTL regions than age-related volume declines, and lead to qualitatively different forms of impairment (note however, that age-related declines in the perforant pathway between the entorhinal cortex and the hippocampus may be present in our sample, see also Yassa, Muftuler, & Stark, 2010). One plausible explanation to reconcile our results with the established literature is that hippocampal processing of object-in-place memory is constrained by its inputs from (or outputs to) the alERC. In the current participant sample, reductions in alERC volume (or alERC hypoactivity in our older adult sample, see Berron et al., 2018 & Reagh et al., 2018) could act as a rate-limiting step for object-in-place memory processes in the hippocampus. Alternatively, the ERC might mediate between the hippocampus and the oculomotor system, interfacing between viewing behavior and representations required for memory (Shen, Bezgin, Selvam, McIntosh, & Ryan, 2016).

Our focus on how alERC volume relates to behavior was inspired by our interest in understanding how alERC cognitive processes might be affected by neurodegeneration. As one of the earliest regions affected by AD, behavioral changes related to alERC volume changes are particularly important as possible indicators of future cognitive impairment. While the cross-sectional nature of this study does not allow us to draw conclusions about whether the differences in visual exploration behavior related to spatial processing of objects we observe is indicative of future cognition, a longitudinal follow-up of this group would answer that question. In the long-term, tasks like the one we present here, perhaps as part of a larger neuropsychological battery of region-specific cognitive tasks, might be informative of dementia-related brain changes. This would be useful both in screening for potential dementia, as well as for assessing patients in places where neuroimaging is expensive or unavailable (as such tasks could be administered remotely; see also Crutcher et al., 2009; Freitas Pereira, Von Zuben A Camargo, Aprahamian, & Forlenza, 2014; Whitehead, Gambino, Richter, & Ryan, 2015; Whitehead, Li, McQuiggan, Gambino, & Ryan, 2018; Zola, Manzanares, Clopton, Lah, & Levey, 2013)

In conclusion, we showed that in older adults, alERC volume was selectively related to viewing behavior towards a “critical object” associated with a scene, but only when the position of that object had been moved within the scene. This study informs a newly evolving understanding of the cognitive role of the alERC, suggesting it may support aspects of spatial processing of objects. Further experimental work will be necessary to elucidate the exact nature of the alERC’s cognitive role, however, this work points to the potential clinical value of eyetracking-based tests for the early detection of dementia.

## Acknowledgements

This work was supported by the Canadian Natural Sciences Engineering Research Council (Discovery and Accelerator grants to M.D.B.; Canada Graduate Scholarship (Doctoral Program) to L.Y.), a Scholar Award from the James S. McDonnell Foundation to M.D.B, and the Canada Research Chairs Program to M.D.B.

## Conflict of Interest

The authors declare no competing financial interests.

